# Bounded Rationality in *C. elegans*

**DOI:** 10.1101/257535

**Authors:** Dror Cohen, Meshi Volovich, Yoav Zeevi, Lilach Elbaum, Kenway Louie, Dino J Levy, Oded Rechavi

## Abstract

Rational choice theory assumes optimality in decision-making. Violations of a basic axiom of economic rationality known as “Independence of Irrelevant Alternatives” (IIA), have been demonstrated in both humans and animals, and could stem from common neuronal constraints. We developed tests for IIA in the nematode *Caenorhabditis elegans*, an animal with only 302 neurons, using olfactory chemotaxis assays. We found that in most cases *C. elegans* make rational decisions. However, by probing multiple neuronal architectures using various choice sets, we show that asymmetric sensation of odor options by the AWC^ON^ neuron can lead to violations of rationality. We further show that genetic manipulations of the asymmetry between the AWC neurons can make the worm rational or irrational. Last, a normalization-based model of value coding and gain control explains how particular neuronal constraints on information coding give rise to irrationality. Thus, we demonstrate that bounded rationality could arise due to basic neuronal constraints.

## Introduction

Humans (Kahneman and Tversky 1979, 1992), and also other animals (Bateson, Healy, and Hurly 2003; Hurly and Oseen 1999; Louie, Khaw, and Glimcher 2013; Royle, Lindström, and Metcalfe 2008; Shafir 1994; Shafir, Waite, and Smith 2002; Yamada et al. 2013), can behave irrationally in many contexts. The neuronal mechanisms that lead to irrational behaviors are still unknown. Failures of “rationality”, i.e inconsistency in preferences, may reflect the implementation of decision-making in biological nervous systems facing intrinsic physical and metabolic constraints (Simon 1955, 1956). Despite varying nervous system architectures, all animal tested behave irrationally, suggesting that rationality and deviations from rationality arise from general computational principles rather than specific biological implementations. According to the idea of *bounded rationality*, the computational or informational load required to make truly optimal decisions exceeds the capacity of our nervous systems (Simon 1955, 1956). One central requirement of rationality and stable value functions is *independence of irrelevant alternatives*, or IIA (Luce. 1959). According to this axiom, a preference between two options should be unaffected by the number or quality of any additional options, and the relative choice ratio between options A and B (pA/pB) should remain constant regardless of the choice set. However, contextual factors such as choice set size significantly alter animal and human decisions.

To examine the boundaries which lead to irrationality, we established *Caenorhabditis elegans* nematodes as a model organism for rational decision making. *C. elegans* has only 302 neurons, 32 of which are chemosensory neurons, and uses chemotaxis to achieve sophisticated behaviors, including simple forms of ethologically-relevant decision making (Barrios 2014; Borne, Kasimatis, and Phillips 2017; Jarrell et al. 2012; Leighton et al. 2014; White et al. 2007). Just two pairs of worm amphid sensory neurons, AWC and AWA, are required for chemotaxis toward attractive volatile odors (Cornelia I. Bargmann, Hartwieg, and Horvitz 1993). Specific odors are known to be sensed exclusively by either the AWC or AWA neurons. The two AWC neurons are structurally similar but functionally distinct from each other, and sense different odors (Bargmann 2006). One detects 2-butanone and Acetone (AWC^ON^), while the other detects 2,3-pentanedione (AWC^OFF^) (Alqadah et al. 2016; Bargmann 2006; Choi et al. 2018; Wes and Bargmann 2001; Worthy et al. 2018).

To investigate if *C. elegans* exhibits non-optimal choice behavior, we conducted odor preference tests, as previously described (Cornelia I Bargmann, Hartwieg, and Horvitz 1993; Ward 1973). To find *“IIA violations”* we measured the relative preference between two attractant spots, in which the most attractive odor A and less attractive odor B were placed, in the presence or in the absence of the least attractive third odor C (Fig.1, A). By changing the concentrations of the odors used in each test, we controlled which specific odorant would be the most attractive (A), the second best (B), and the least attractive option (C).

**Fig. 1.**
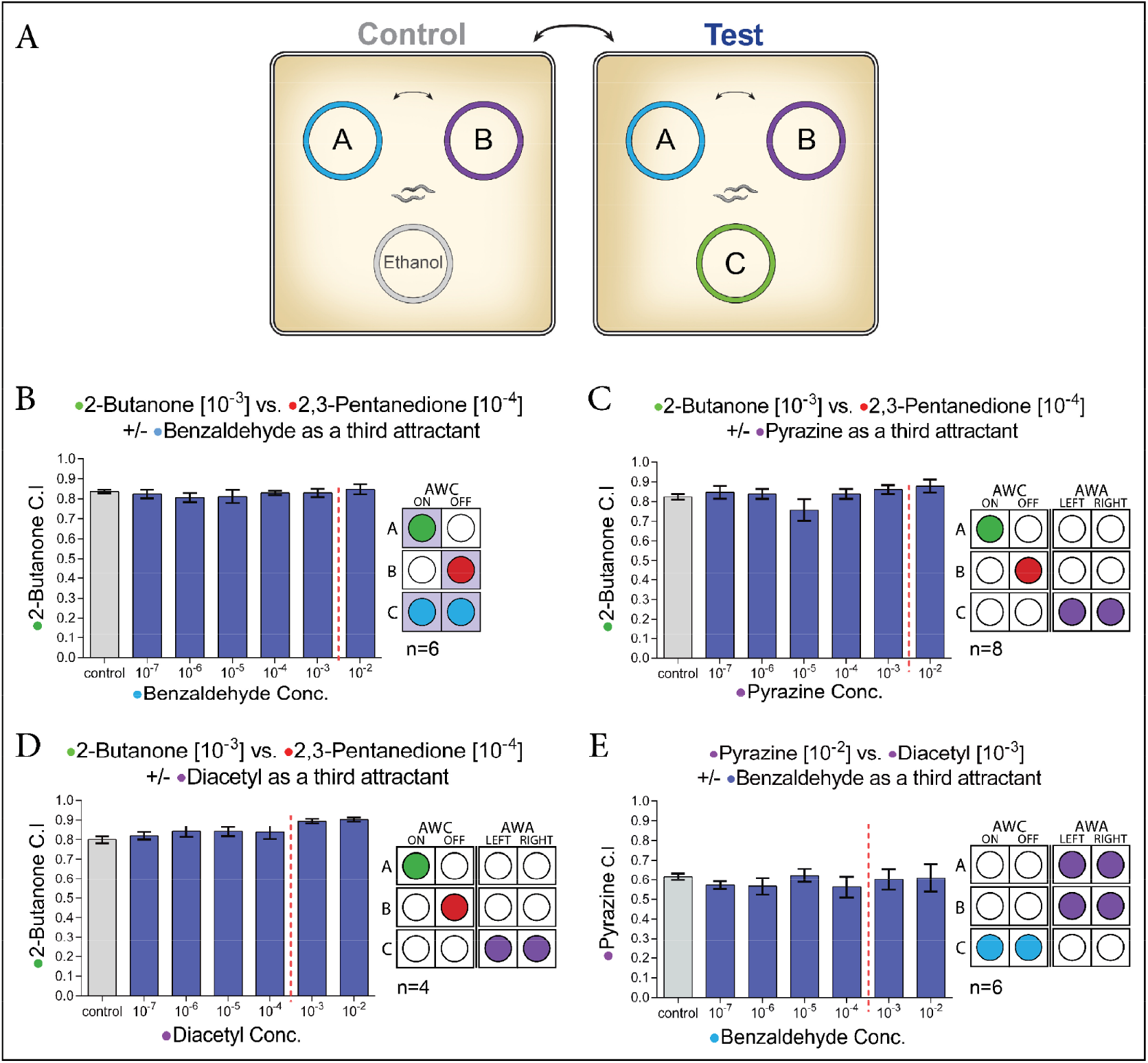
*C. elegans* display rational decisions. **(A)** A scheme for the Independence of Irrelevant Alternatives chemotaxis assays. A chemotaxis index **(C.I)** (number of worms in A, divided by the number of worms in A and B together) was calculated. Each plate contained ~200-400 worms. **(B)** The relative preference for 2-butanone over 2,3-pentanedion is unaffected by increasing concentration of benzaldehyde as a third attractant (n=6). **(C-D)** introducing AWA sensed odorants as a third attractant, does not influence the relative preference between 2-butanone and 2,3-pentanedione (C: n=4, D: n=8). **(E)** benzaldehyde as a third attractant, does not affect the relative preference between the two AWA sensed odorants pyrazine and diacetyl (n=6). Bars represent the C.I of odor A. Dashed red lines indicate the point where odor C was too attractive for our purposes, i.e. it rendered B irrelevant to the choice task. Colors correlate between an odor and the specific neuron recognizing it. Wilcoxon Signed-Ranks Test, Error bars represent the standard error of the mean C.I.

We first used odors which are sensed by a minimal decision-making neuronal circuit, by performing choice assays with odors detected exclusively by the two AWC neurons: 2-butanone (odor A), 2,3-pentanedione (odor B), and benzaldehyde (odor C) (Fig.1, B and **Fig.S1**). In these experiments the third odor C was sensed by the neurons that sense both odor A and odor B, in a balanced/symmetric way. Namely, since odor C was sensed by both AWC neurons, it could potentially disrupt the sensing of both odor A (sensed only by the AWC^ON^ neuron) and odor B (sensed only by the AWC^OFF^ neuron). We tested the effect that different concentrations of odor C would have on the relative preference between A and B. While IIA violations have been observed in a wide variety of organisms (Bateson et al. 2003; Hurly and Oseen 1999; Louie et al. 2013; Royle et al. 2008; Shafir 1994; Shafir et al. 2002; Yamada et al. 2013), despite numerous repetitions and iterations, we found that in *C. elegans* the addition of increasing concentrations of odor C did not lead to violations of rationality, as the preference ratio between odor A and odor B did not change in a statistically significant or physiologically relevant way. Specifically, in no case did odor B become more attractive relatively to A (Fig.1, B and **Fig.S1**).

In many organisms the dopaminergic system has a strong effect on decision-making (Doya 2008; Rogers 2011). Therefore, we subjected *cat-2* mutants, defective in dopamine synthesis, to the same choice task described above. Similarly to wild type animals, we did not observe any statistically significant differences between wild-type and *cat-2* mutants in the preference between odors A and B in the presence or in the absence of option C (**Fig.S2**). These experiments suggest that in *C. elegans* the lack of dopamine signaling does not lead to IIA violations.

In the experiments described above, all three odorants (A, B, and C) were sensed by just two neurons, AWC^ON^ and AWC^OFF^. It is possible that this minimal neuronal circuit was “too simple” to give rise to inconsistent behaviors - perhaps irrationality stems from complexity? To increase the complexity of the neuronal circuit underlying the decision process, we tested combination of odors that are sensed by both AWC and AWA neurons. We started by testing “balanced” third odors C, in the sense that these odors are not sensed preferentially just by the neurons that sense odor A or odor B. We found that increasing the circuit complexity through the addition of another pair of neurons does not lead to inconsistencies in decision-making (Fig.1, C-E). All the results presented above demonstrate that the worm’s decision-making process can be consistent and robust at least when the irrelevant alternative is sensed symmetrically, in a balanced way, by the neurons that sense odor A, and the neurons that sense odor B.

Next, we broke the symmetrical pattern of olfactory inputs, to test if asymmetry in the sensing of the different odors can lead to irrational decision making. In a number of experiments testing different sets of odors, we found that IIA violations can occur due to an *asymmetric overlap* between odors A and C, independently of the number of neurons involved (Fig.2, A-B). More specifically, we found that IIA violations can occur when odor C is sensed in an imbalanced manner by the neurons that sense odor A but not odor B. To our knowledge, this is the first demonstration of economic irrationality and IIA violations in *C. elegans*.

**Fig. 2.**
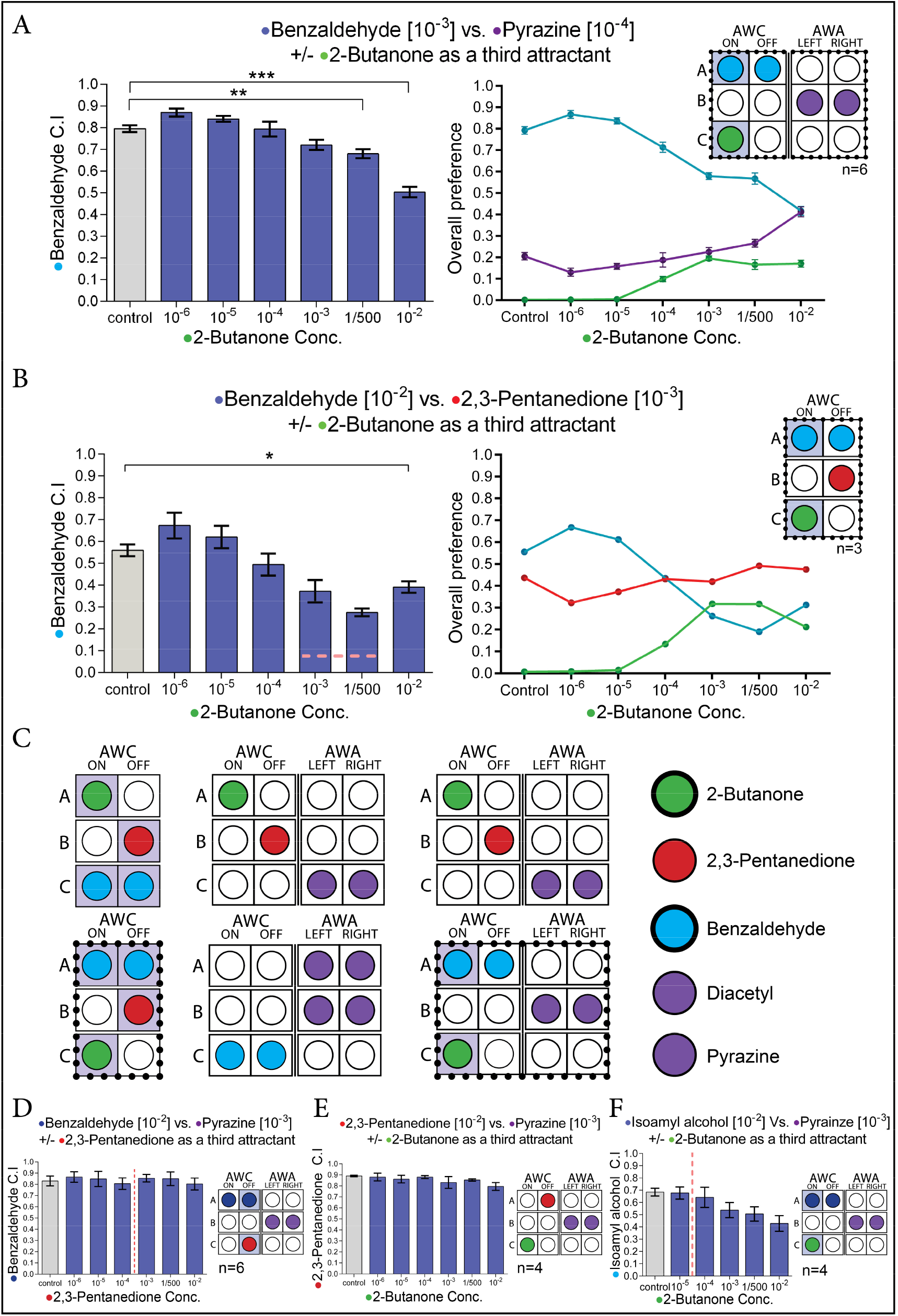
*C. elegans* exhibit IIA violations when specific neuronal architectures are induced. **(A)** The effect of 2-butanone as a third attractant on the relative preference between benzaldehyde and pyrazine, and the overall preference of each attractant point in every condition (Wilcoxon Signed-Ranks Test, C=1/500: W=5, P<0.003 ; C=10^-2^: W=0, P<0.000 ; n=6). **(B)** The effect of 2-butanone as a third attractant on the relative preference between benzaldehyde and 2,3-pentanedion, and the overall preference of each attractant point in every condition. (Wilcoxon Signed-Ranks Test, C=10^-2^: W=0, P<0.023 ; n=3). **(C)** In all the violations that we described so far, 2-butanone, sensed specifically by the AWC^ON^ neuron, functioned as odor C, and benzaldehyde, sensed by both AWC neurons, functioned as odor A. **(D)** 2,3-pentanedione as a third attractant does not change the relative preference between benzaldehyde and pyrazine (Wilcoxon Signed-Ranks Test, n=6). **(E)** 2-butanone as a third attractant does not change the relative preference between 2,3-pentanedione and pyrazine (Wilcoxon Signed-Ranks Test, n=4). **(F)** 2-butanone as a third attractant significantly reduced the relative preference for isoamyl-alcohol over pyrazine (Wilcoxon Signed-Ranks Test, C=1/500: W=3, P<0.042 ; C=10^-2^: W=1, P<0.012 ; n=4). Bars represent the C.I of odor A. Dashed red lines indicate the point where odor C was too attractive for our purposes, i.e. it rendered B irrelevant to the choice task. Colors correlate between an odor and the specific neuron recognizing it. Error bars represent the standard error of the mean C.I.

In all the violations that we described so far, 2-butanone, sensed specifically by the AWC^ON^ neuron, functioned as odor C, and benzaldehyde, sensed by both AWC neurons, functioned as odor A. Thus, an alternative explanation to these results is that the violations do not occur from breaking of the symmetry in the odors’ sensation, but arise instead from a specific interaction between butanone (C) and benzaldehyde (A) (Fig.2, C). When 2,3-pentanedione (an odor which is sensed by AWC^OFF^) served as odor C instead of 2-butanone, the worms behaved rationally (Fig.2, D). It was previously reported that the asymmetry between the two AWC neurons (having both AWC^ON^ and AWC^OFF^ neurons) is required for the ability to discriminate between benzaldehyde and 2-butanone. The authors hypothesized that 2-butanone can attenuate benzaldehyde signaling in the AWC^ON^ neuron (Wes & Bargmann, 2001). Therefore, we conducted different experiments to test if the observed IIA violations stem from specific interactions between 2-butanone and benzaldehyde, in the AWC^ON^ neuron. Four lines of evidence suggest that this is not the case, and instead, a general circuit principle of asymmetry in sensation underlies irrationality.

First, when 2-butanone served as odor A, and benzaldehyde served as odor C, no violations were observed (see Fig.1, B). Thus, the worms do not make inconsistent decisions simply because they cannot distinguish between these two odors.

Second, using 2-butanone as odor C is not enough to make the worms irrational; when 2-butanone serves as odor C, but the circuit was symmetrical, the worm made consistent, rational decisions and did not show IIA violations (Fig.2, E). This shows that 2-butanone cannot be considered as a general “distractor” or “confusant” molecule, similarly to the repellent DEET pesticide (Dennis et al. 2017).

Third, while both benzaldehyde and 2-butanone are attractive odors when presented separately (Cornelia I. Bargmann et al. 1993), little is known about the ecology of *C. elegans* (Cornelia I. Bargmann et al. 1993; Sagasti et al. 1999; Schulenburg and Félix 2017), and it is possible that the combination of the two odors in the wild is associated with an unattractive or even repulsive substance. In this case it would be rational for the worm to avoid an unattractive odor, formed from the combination of benzaldehyde and 2-butanone, when both are present on the same plate - it would be a “feature”, not a “bug. To test this possibility, we examined if worms prefer benzaldehyde over a mixed combination of benzaldehyde and 2-butanone (see Materials and Methods). We found that the combination of 2-butanone and benzaldehyde was more attractive than benzaldehyde alone. The spot that contained both odors was as attractive as would be expected based on the simple summation of the attractiveness of each of the odors alone (**Fig.S3**). Thus, introducing 2-butanone on to the plate does not create a new unattractive odor (with benzaldehyde) that can explain the IIA violation that we observed. These results strengthened the hypothesis that the IIA violations that we observed are due to constraints on the neural system – that is, it’s a “bug”, not a “feature”.

Fourth, we found that IIA violations arise also due to exposure to other odors, not just benzaldehyde or butanone, when odor C is sensed asymmetrically: by the neurons that sense odor A, but not by the neurons that sense odor B. When isoamyl alcohol was used as odor A instead of benzaldehyde (the two chemicals are sensed by both AWC neurons), the worms behaved inconsistently (when odor C was sensed asymmetrically) (Fig.2, F). Further, when acetone was used as odor C instead of butanone (both are sensed only by the AWC^ON^ neurons) the preference towards odor A was also disproportionally reduced (**Fig.S4**).

In summary, the violations of rationality that we documented do not arise exclusively because of the two odorants benzaldehyde and butanone, but stem from a general property of the asymmetry in the sensation of the odor choices.

The asymmetry that we found to drive IIA violations, led us to hypothesize that the AWC^ON^ neuron has special limitations that make it uniquely susceptible towards inconsistent choice behavior. We therefore tested if mutants which have two AWC^ON^ neurons (AWC^ON/ON^, loss of AWC asymmetry), would be more prone to inconsistent decision making. When the AWC^ON^-sensed odor acetone was used as odor C, AWC^ON/ON^ mutants made more inconsistent decisions in comparison to wild type worms, and unlike wild type worms, even showed a preference reversal between odors A and B (Fig.3, A and **Fig.S4**). As with acetone, the “distracting” effect of 2-butanone, when used as odor C, was much stronger in AWC^ON/ON^ mutants compared to wild type (in comparison to wild type, the preference reversal occurred in lower concentrations of odor C) (Fig.3, B). Note, that the AWC^ON/ON^ mutants are hypersensitive to acetone and butanone, since they sense these odors using two AWC^ON^ neurons instead of one; therefore in relatively low concentrations, these odors instantly became the most attractive odors on the plate (**Fig.S5**). For this reason, while we detected preference reversals with the AWC^on/on^ mutants, the effects cannot be considered *bone fide* IIA violations.

**Fig. 3.**
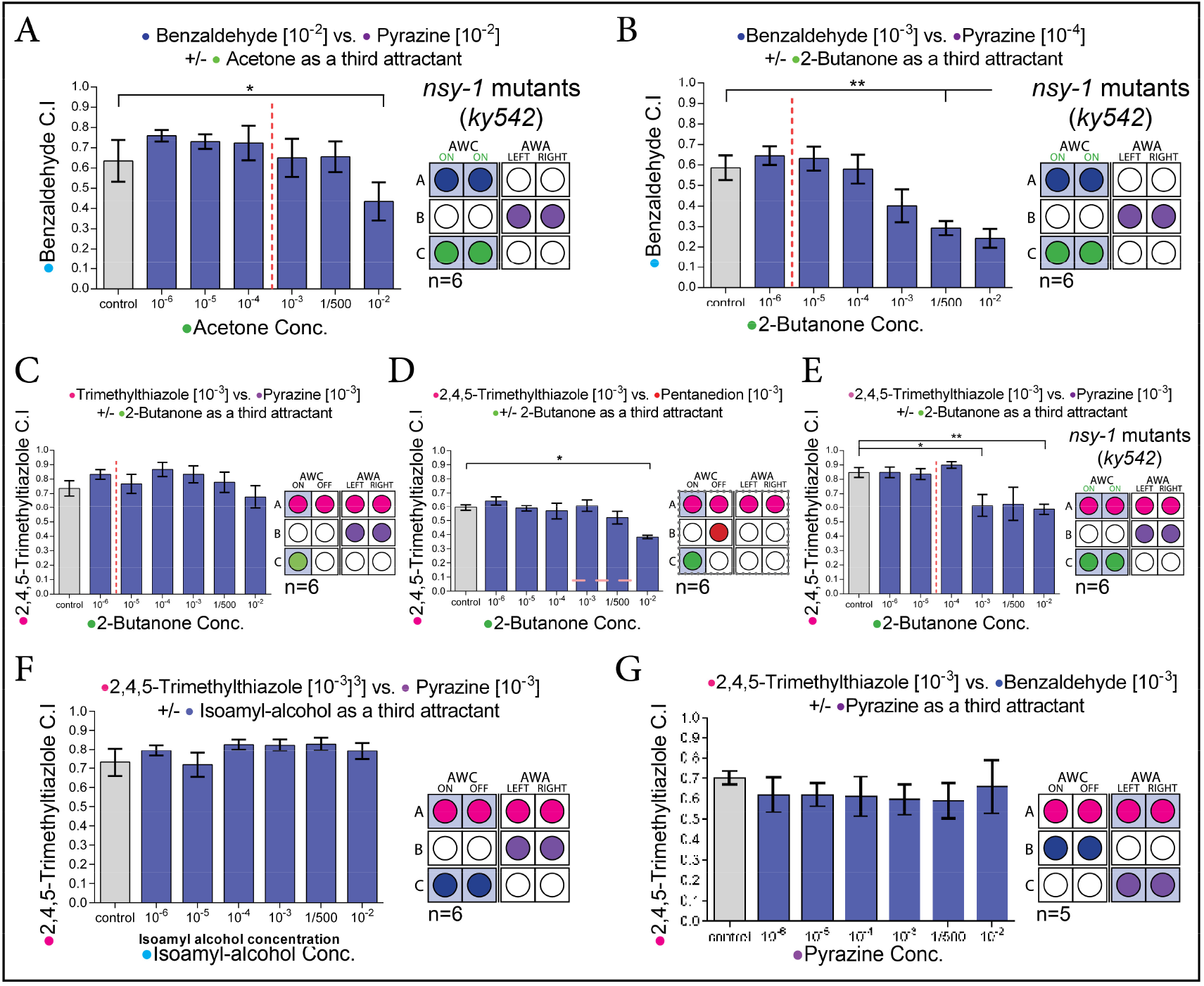
The AWC^on^ neuron makes the worm vulnerable to IIA violations. **(A)** The influence of acetone as a third attractant on the relative preference between benzaldehyde and pyrazine, in AWC^ON/ON^ mutant worms. (Wilcoxon Signed-Ranks Test, C=10^-2^: W=13, p=0.014; n=6). **(B)** The influence of 2-butanone as a third attractant on the relative preference between benzaldehyde and pyrazine, in AWC^ON/ON^ mutant worms. (Wilcoxon Signed-Ranks Test, C=1/500: W=2, p=0.004; C=10’^2^: W=2, p=0.004; C=1/500: W=2, p=0.004; n=6). **(C)** 2-butanone as a third attractant does not change the relative preference between 2,4,5-trimethylthiazole and pyrazine (Wilcoxon Signed-Ranks Test, n=6). **(D)** The influence of 2-butanone as a third attractant on the relative preference between 2,4,5-trimethylthiazole and 2,3-pentanedione (Wilcoxon Signed-Ranks Test, C=10^-2^: W=0, P<0.004 ; n=6). **(E)** 2-butanone as a third attractant change the relative preference between 2,4,5-trimethylthiazole and pyrazine in AWC^ON/ON^ mutant worms (C=10^−2^: W=0, p=0.002; C=1/500: W=7, p=0.093; C=10^−3^: W=4, p=0.025; n=6). **(F)** Isoamyl-alcohol as a third attractant does not change the relative preference between 2,4,5-trimethylthiazole and pyrazine (Wilcoxon Signed-Ranks Test, n=6). **(G)** Pyrazine as a third attractant does not change the relative preference between 2,4,5-trimethylthiazole and benzaldehyde (Wilcoxon Signed-Ranks Test, n=5). Bars represent the C.I of odor A. Dashed red lines indicate the point where odor C was too attractive for our purposes, i.e. it rendered B irrelevant to the choice task. Colors correlate between an odor and the specific neuron recognizing it. Error bars represent the standard error of the mean C.I.

We continued to study how constraints in AWC^ON^ sensation bound rationality. It is possible that irrationality arises due to interference of odor C with the sensing of odor A in the AWC^ON^ neuron. As noted above, in AWC^ON/ON^ mutants, 2-butanone interferes with the sensing of benzaldehyde in the AWC^ON^ neuron (Wes and Bargmann 2001). We examined whether acetone, which like 2-butanone induces irrationality and is sensed only by the AWC^ON^ neuron, also disturbs the sensing of benzaldehyde in the AWC^ON^ neuron. Indeed, we found that in AWC^ON/ON^ mutants, acetone interferes with benzaldehyde sensation (**Fig.S6**). Together, all these experiments strengthen the case for AWC^ON^ being uniquely vulnerable for sensation of multiple different competing odors, a limitation which in certain cases can lead to IIA violations in olfactory choice behaviors.

Can minimizing the role of the AWC^ON^ neuron in the sensation of odor A expand the boundaries of rationality? To test this we diluted the role of AWC^ON^ in sensation of odor A, and examined if this “buffers” against AWC^ON^-dependent irrationality. As odor A, we switched benzaldehyde (sensed by the AWC neurons) with 2,4,5-trimethylthiazole, which is sensed by both the AWC and AWA neurons (Bargmann 2006; Cornelia I Bargmann et al. 1993). We examined two odor setups, and when the role of AWC^ON^ in the sensation of odor “A” was reduced, we did not observe consistent IIA violations (Fig.3, CD). Further, in AWC^ON/ON^ mutants, where the role of the AWC^ON^ neuron is increased, we did find strong changes in the preference ratio of A Vs. B (Fig.3, E). These experiments suggest that the relative weight of the AWC^ON^ in the sensation of odor A affects the tendency to demonstrate inconsistent behavior.

We tested also if the relative role of AWC^ON^ in sensation of odor C, the agent of confusion, makes a difference. In additional experiments, we “expanded” the circuit, to dilute the role of AWC^ON^ in sensation of odor C, while preserving the proportion of neurons that sense each odor. Each of the odors was sensed by twice as many neurons (in comparison to the experiment described in Fig.2, B). In these experiments, when odor C was sensed also by the AWC^OFF^ neuron, we did not observe any IIA violations, nor did we see any significant changes in the preference of A over B (Fig.3, F). Similarly, when odor C was sensed by the AWA neurons we did not observe any IIA violations, nor did we see any significant changes in the preference of A over B (Fig.3, G). Thus, the relative weight of the AWC^ON^ in the sensation of both odors A and C affects the capacity of the worm to behave rationally.

Given our behavioral results, we propose a model of pathway-specific sensory gain control and examine its predictions (Fig.4). The essential feature is that (at least some) neurons in the chemosensory pathway perform a form of sensory gain control analogous to divisive normalization (see Materials and Methods). Critically, cross-odorant gain control only occurs when a given neuron is sensitive to more than one odor and both those odors are present at the same time (i.e. in the same choice set). Specifically, increasing concentrations of odor C will divisively scale responses to odor A (when A and B are fixed). Odor B representations, being independent from odor C coding, are unaffected by concentrations of C. Thus, the general prediction is that increasing C will decrease the relative preference of A over B.

**Figure 4.**
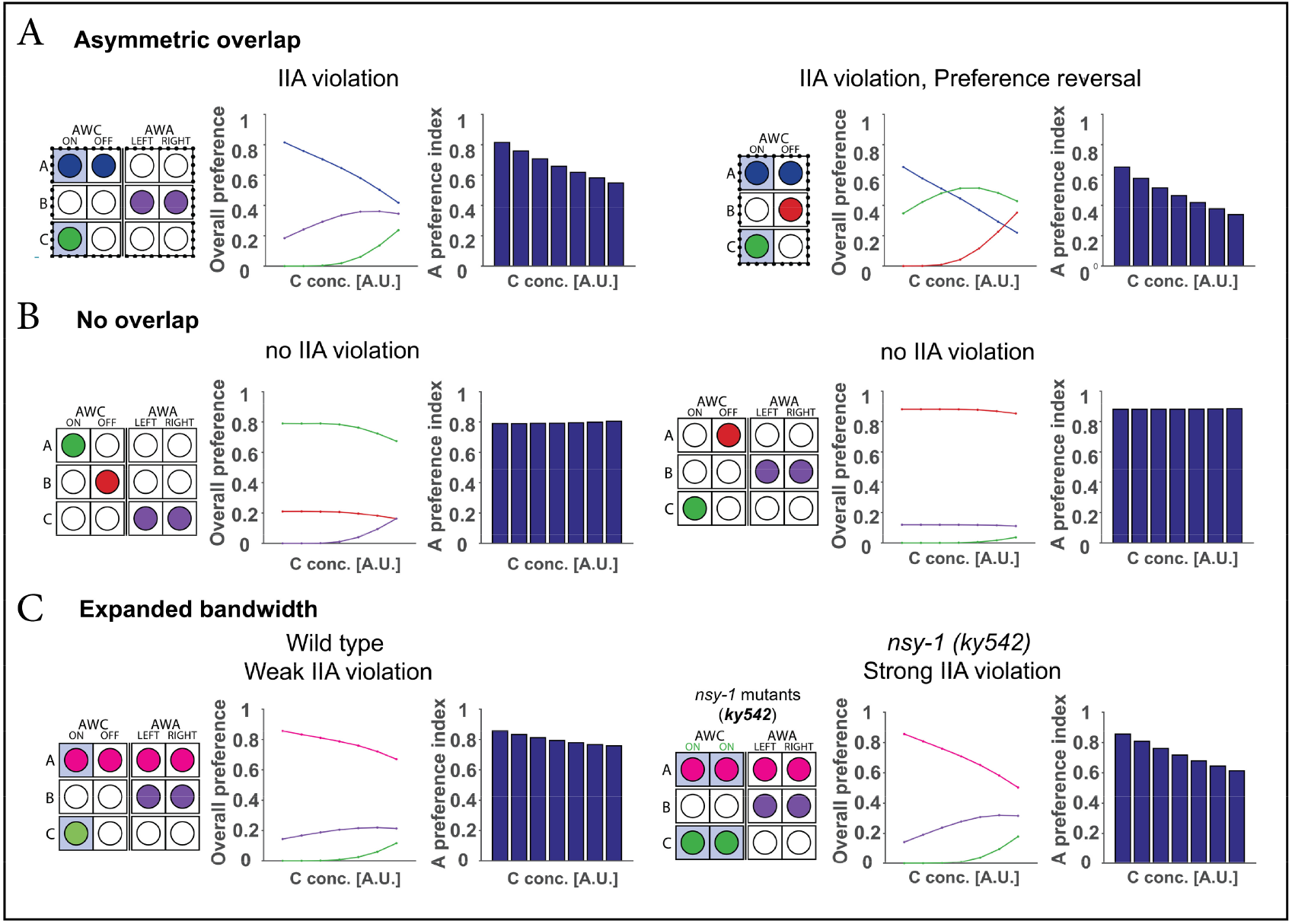
Sensory gain control model of chemosensation explains circuit architecture-specific IIA violations. Predicted choice behavior in a divisive normalization model of sensory gain control in *C. elegans* chemosensation. Odors driving the same chemosensory neuron in a choice scenario are assumed to drive cross-odor normalization in neural representation; specifically, this cross normalization is assumed to be stronger in AWC^ON^ neurons. Simulation parameters were not fit to empirical choice data, but instead were chosen to demonstrate qualitative similarity in behavioral data under different circuit activation patterns. Each trinary scenario was simulated for *n*=10^6^ repetitions. In each trinary combination shown below, the left panel shows the circuit activation pattern, the middle panel shows model-precited preference for the three odors at different concentraitons of odor C, and the right panel shows the preference index for odor A (relative choice of odor A vs. odor B). Our results indicate that IIA violations can occur due to an *asymmetric overlap* between odors “A” and “C”. **(A)** Model-predicted IIA violations in asymmetric overlap circuit architectures. When odor A and odor C both activate AWC^ON^, increasing concentrations of odor C reduce the representation of odor A in the model via cross-odor normalization. Note that the model can capture both IIA violations without preference reversals (left) and IIA violations with preference reversals (right). Both types of IIA violations are observed in the emiprical choice data. **(B)** Model-predicted rational choice behavior (no IIA violations) in non-overlap circuit architectures. In these circuits, odor A and odor C activate distinct chemosensory neurons and no cross-odor normalization occurs in the model. Thus, the neural representations of odors A and B (and the relative choice preference of A over B) do not vary with the concentration of odor C. **(C)** Model behavior in expanded bandwidth circuits. The model assumes that cross-odor normalization occurs only (or most strongly) in AWC^ON^ neurons. In expanded bandwidth scenarios, odor A activates both AWC and both AWA neurons. In wild-type worms, cross-odor normalization only affects 25% of the neural representation of odor A and the model predicts weak IIA violations (left). In *nsy-1 (ky542)* mutants with two AWC^ON^ neurons, cross-odor normalization affects 50% of the neural representation of odor A and the model predicts stronger IIA violations (right).

Our simulations show that the model predicts empirically observed IIA violations in asymmetric overlap scenarios (Fig.4, A), capturing the decrease in relative preference of A vs. B as C increases. Furthermore, the model can also capture two additional aspects of observed choice (Fig.4, A**, right**): (1) preference reversal of odors A and B, and (2) eventual selection of odor C over odor A. The gain control model also explains why *C. elegans* display rational choice in other circuit activation scenarios (Fig.4, B), where odor C is sensed in different chemosensory neurons than those sensing odors A and B. In the model, this translates into the equations for *R_A_* and *R_B_* carrying no C-related terms in the divisive denominator: the activity representing A and B - and thus the relative preference between the two – is independent of distractor odor C.

With additional circuit-specific clarification, the gain control model explains observed choice behavior in expanded bandwidth scenarios, where the sensing of odor A involves all four chemosensory neurons (Fig.3). Model predictions for both wild-type and AWC^ON/ON^ mutants emphasize, as the empirical data suggest, a particularly important role for the AWC^ON^ neuron in mediating cross-odor gain control (Fig.4, C). The expanded asymmetric scenario is functionally similar to the simpler asymmetric overlap scenario in which IIA violations were observed. However, despite this analogous organization, the expanded scenario does not generate IIA violations. Why might this be the case? The gain control model captures this IIA consistency by positing that cross-odor gain control is specific to (or stronger in) AWC^ON^ neurons. Thus, an odor (e.g. 2,4,5-trimethylthiazole) sensed by all four chemosensory neurons exhibits gain control effects in only 25% of its representation, while an odor sensed by only AWC^ON^ and AWC^OFF^ (e.g. benzaldehyde) exhibits gain control in 50% of its representation; the model thus predicts a diminished effect of contextual odors on choice behavior in expanded bandwidth scenarios. Furthermore, consistent with the data, the model predicts that AWC^ON/ON^ mutants-which exhibit the equivalent of two functional AWC^ON^ neurons – should exhibit stronger IIA violations than wild-type worms in identical choice conditions.

In this work we demonstrate for the first time that even *C. elegans*, with its extremely minimal nervous system, displays IIA violations in decision making. In most cases worms behave rationally, however, IIA violations do occur when the different options are represented in an imbalanced way in the AWC^on^ neuron.

We suggest that the evolution of the AWC^ON^ neuron, which expanded the repertoire of odors that *C. elegans* can sense, came at the cost of inconsistency in decision making. This demonstrates a simple example of the phenomenon known as the “paradox of choice” (Iyengar and Lepper 2000; Schwartz 2003), where more options are normatively considered better but in many cases, lead to suboptimal choices.

Our experimental results and normalization-based mathematical model are aligned with previous studies on cortical computation which showed that IIA violations can be explained by a divisive normalization framework (Louie, Grattan, and Glimcher 2011; Louie et al. 2013; Webb, Glimcher, and Louie 2014). Thus, our work supports the notion that IIA violations are a result of neural constraints carved by evolution to maximize information under limited time and resources (Cochella et al. 2014; Palmer 1996), and directly relate to the high-level concept of “bounded rationality” (Simon 1972). We propose that understanding the building blocks of choice in an animal with a compact, deciphered, rigid, and stereotypic connectome, can shed light on the fundamental biological constraints and principles that generate (non)-rational behavior in simple as well as in complex organisms.

## Materials and methods

### Strains and husbandry

The strains used in this work: Bristol N2 wild-type and *cat-2* (n4547) and nsy-1(ky542). Bristol N2 wild-type and *cat-2* (n4547) strains were provided by the CGC, which is funded by NIH Office of Research Infrastructure Programs (P40 OD010440). All strains were maintained at 20°C on NGM plates supplemented with the antifungal agent Nystatin and fed with *E. coli* OP50 (Stiernagle 2006).

### Obtaining synchronized worms (“Egg-prep”)

A synchronized population of worms was obtained by employing a standard “egg-prep” procedure, as previously described (Stiernagle 2006).

### Chemotaxis Assays

Chemotaxis assays were based on classical chemotaxis assays, (Cornelia I. Bargmann et al. 1993; Ward 1973). Assay plates were square 12X12cm dishes containing 30 ml of 1.6% BBL agar (Benton-Dickinson), 5mM potassium phosphate (pH 6.0), 1mM CaCI2 and 1mM MgSO4. Three marks were made on the back of the plates equidistant from the center of the plate (3cm) and from each other (5.2cm). The diluted attractants (1 μl) was placed on the agar over one marks. In the control plates (binary choice), 1 μl of 100% ethanol was placed over the third mark (all attractants were diluted in ethanol). The tested animals were placed at the center of the plate, equidistant from the three marks. Attractants were obtained from Sigma-Aldrich. Pure pyrazine is a solid, so pyrazine dilutions are weight:volume rather than volume:volume as for other attractants.

Well-fed adult animals were washed three times with wash buffer (0.5% Gelatin, 5mM potassium phosphate (pH 6.0), 1mM CaCI2 and 1mM MgSO4), then placed near the center of a plate equidistant from the attractants (and the control spot when present). Approximately one hour after the assay began, the numbers of animals at the three areas (2 cm radius of each attractant) were determined, as well as the total number of animals in the assay, the number of animals that were not at any attractant area, and the number of animals that stayed in the starting point (did not cross a 1cm diameter circle around the center of the plate). A specific C.I was calculated as:

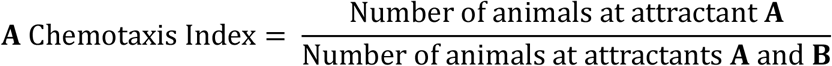

The C.I could vary from 0 to 1. The animals were anesthetized when they reached the attractant. 1μl of Sodium azide 1M was placed at each one of the three spots, 15 minutes in advanced. Sodium azide anesthetized animals within about a 1 cm radius of the attractant. For discrimination assays, aceton was added to a final concentration of 1.2 μl per 10-ml plate, and mixed with the liquid agar once it had cooled to 55 °C.

### “Bug” or “Feature” assays

We measured the relative preference between 2ul of benzaldehyde (10^-2^) (A) and 2ul of butanone (10^-2^) (B), and compared it to the relative preference between 2ul of benzaldehyde (10^-2^) (A), and a mixture of 1ul of butanone (1/50) and 1ul of benzaldehyde (1/50) (A’B’). The butanone spot (B) and the butanone+benzaldehyde (A’B’) spot, contain the same amount of butanone molecules, as well as an equal volume of ethanol. The “A’B’” spot contains, in addition to butanone, the same amount of benzaldehyde molecules as presented by “A”. Each assay included 3 “A vs. B” plates, coupled to 3 “A vs. A’B’” plates. Each data point represents the mean of 6 essays performed on two different days.

### Statistical analysis

Data are presented as mean +/− SEM. Statistical significance of differences in chemotaxis index between control and test plates in a certain concentration were analyzed by **A Wilcoxon Signed-Ranks Test** (*P*<0.05 was regarded as significant; “*” means *p*< 0.05, “**” means *P*<0.01, “***” means *p*< 0.001, and “****” means *P*<0.0001.)

### Normalization model of sensory gain control

To examine whether both IIA and non-IIA choice behavior can be explained by a circuit-specific model of sensory gain control in chemosensation, we implemented a simple divisive normalization-based computaitonal model of chemosensory value coding. Gain control is a widespread representational principle in early sensory processing in which the overall level of coding activity is regulated by the specific context present at the time of encoding. For example, Drosophila antennal lobe neural activity representing a specific odor will depend on whether other odors are present, and primate primary visual cortical responses to a center stimulus will be suppressed by stimuli in the sensory surround. Many of these gain control interactions can be explained a normalization computation, in which the feedforward-driven response of a neuron is divided by a term that represents a larger pool of neurons. This normalization pool (acting via the equation denominator) provides a mechanism for contextual modulation of the stimulus-specific response (in the numerator). For example, the response of Drosophila antennal lobe neurons are increased by a test odorant but suppressed by a mask odorant, a pattern described by normalization-based gain control.

To examine the predictions of a gain control model of chemosensation, we constructed a model of pathway-specific sensory gain control in *C. elegans* chemosensation and explored its qualitative predictions in trinary odorant choice behavior. In this model, neural activity *R_i_* representing the decision value of an odorant stimulus *i* depends on its concentration (or intensity) *I_i_* via a divisive normalization representation:

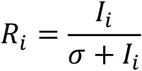

where the semisaturation term σ controls how the function approaches saturation. Context-dependence in this model is instantiated as cross-odorant gain control when a given chemosensory neuron responds to more than one odor and both odors are present in the choice set. For example, in the most basic version of this model, the responses to two odors A and B will be described by the equations:

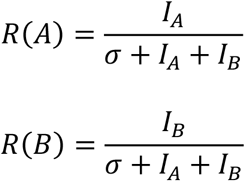

where *I_A_* and *I_B_* are properties of the odor stimuli (i.e. concentrations) and the responses *R*(*A*) and *R*(*B*) denote neural activity representing the value of the odors. Note that this is an algorithmic model intended to model information processing rather than biophysical implementation; however, because *C. elegans* neurons are generally thought to not exhibit action potentials, *R* can be viewed as a graded voltage signal. Furthermore, since this activity integrates across chemosensory neurons, it represents information at a downstream stage: synaptic input to interneurons, interneuron activity, or a more global measure of preference (e.g. turn/run balance in the klinokinesis-governing circuit).

In this model, odorants represented by multiple chemosensory neurons (e.g. benzaldehyde activating both AWC^ON^ and AWC^OFF^) receive a weighted averaging across the active neurons. For example, the response to benzaldehyde (denoted A here) in the presence of butanone (AWC^ON^ only, here denoted B) is described as:

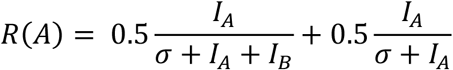

where there is an equal weighting of the responses of AWC^ON^ (left term) and AWC^OFF^ (right term). In general, we assume equal weighting of all chemosensory neurons contributing to a representation, though we relax this when considering the results from odors with expanded bandwidth (four neuron) representations (see below). Note that for simplicity and parsimony, we assume that neurons that are encoding a single odor at a given time (AWC^OFF^ in the example above) also have an analogous form of gain control over the single represented odor. Decisions are implemented by a simple noisy decision rule, assuming a fixed Gaussian noise term (equal across all options in a choice set).

## Acknowledgments

We thank all the Rechavi lab and Levy lab members for helpful discussions. D.C and M.V wish to thank the Sagol School of Neuroscience. We thank Cornelia Bargmann for sharing with us the *nsy-1*(ky542) mutants. We thank Yoav Benjamini, supervisor of Yoav Zeevi, for statistical advice. O.R is thankful to the Adelis foundation grant #0604916191 and ERC grant #335624for funding. D.J.L is thankful to the ISF grant #1104/13 and to the Henry Crown Institute of Business Research for funding. K.L. is thankful to NIH grant R01MH104251 for funding.

